# Construction of a genome-wide pooled CRISPRi library as a resource for exploring the acid tolerance mechanism in *Streptococcus mutans*

**DOI:** 10.1101/2025.07.22.666072

**Authors:** Yaqi Chi, Yuxing Chen, Chongyang Yuan, Liuchang Yang, Mingrui Zhang, Xiaolin Chen, Yiran Zhao, Ming Li, Xiaoyan Wang, Yongliang Li

**Affiliations:** Department of Cariology and Endodontology, Peking University School and Hospital of Stomatology & National Center for Stomatology & National Clinical Research Center for Oral Diseases & National Engineering Research Center of Oral Biomaterials and Digital Medical Devices & Central Laboratory, Beijing, PR China; School of Stomatology, Hebei Medical University, Shijiazhuang, PR China; CAS Key Laboratory of Microbial Physiological and Metabolic Engineering, State Key Laboratory of Microbial Resources, Institute of Microbiology, Chinese Academy of Sciences, Beijing, China; College of Life Science, University of Chinese Academy of Sciences, Beijing, China

**Keywords:** *Streptococcus mutans*, pooled CRISPRi library, next-generation sequencing, acid-tolerance

## Abstract

*Streptococcus mutans* is recognized as the primary etiological agent of dental caries, one of the most prevalent infectious diseases globally. Its remarkable acid tolerance enables survival and proliferation in the low-pH biofilm microenvironment, establishing *S. mutans* as the dominant species in dental plaque and a key contributor to cariogenesis. Although numerous studies have identified genes linked to acid tolerance mechanisms, the full set of essential acid tolerance genes within its genome remains incompletely characterized, largely due to the lack of systematic, genome-scale investigations. To address this knowledge gap, we constructed a genome-wide pooled CRISPR interference (CRISPRi) library targeting 95% of the predicted *S. mutans* genes and employed next-generation sequencing to identify acid tolerance determinants systematically. Our screen revealed 95 acid tolerance-associated genes, a subset of which were functionally validated through gene knockout studies. Functional enrichment analysis demonstrated significant associations with metabolic pathways (including cofactor biosynthesis and amino/nucleotide sugar metabolism), tRNA modification, and transcriptional regulation. Protein-protein interaction (PPI) network analysis identified critical interactors (ComYC, SMU_1979c, DeoC, AcpP, NadD, and SMU_1988c) and two functionally cohesive modules. These findings provide novel mechanistic insights into the acid adaptation strategies of *S. mutans* and highlight potential therapeutic targets for caries prevention.

## Introduction

Dental caries is a common disease in children and adults, affecting approximately 27,500 per 100,000 population worldwide with untreated cases. Its high prevalence and low treatment rates pose significant public health challenges and exacerbate socioeconomic burdens^[1]^. The oral microbiota is closely associated with the initiation and progression of dental caries, and the acid-producing and acid-tolerant bacterial species are recognized as primary pathogens in cariogenesis^[2]^. Among these, *Streptococcus mutans* (*S. mutans*) serves as a hallmark representative. *S. mutans* can rapidly ferment various carbohydrates and produce a large amount of organic acids, decreasing the local pH, ultimately resulting in demineralization of enamel and caries development. *S. mutans* could decrease pH to levels that are toxic to some microbes and produce exopolysaccharides (EPS), creating an acidic niche that promotes acid-tolerant biofilm formation^[3]^.

To survive acid stress, *S. mutans* has evolved multiple physiological adaptation mechanisms, which are collectively referred to as the acid tolerance response (ATR)^[4]^. The F-ATPase in *S. mutans* hydrolyzes ATP to actively extrude H^+^ from the cell^[5,6]^. Notably, its F-ATPase has a lower optimal pH value than that of many other oral microorganisms^[7]^. Studies demonstrate that unsaturated fatty acids (UFAs) in cytoplasmic membranes are critical for acid tolerance^[8]^, as genetic disruption of the *fabM* gene (encoding a key enzyme for UFA biosynthesis) impairs unsaturated fatty acid production, consequently attenuating the mutant strain’s cariogenicity^[9]^. The agmatine deiminase system (AgDS) catabolizes arginine to generate amines, CO₂, NH₃, and ATP, collectively facilitating the maintenance of the transmembrane pH gradient (ΔpH)^[10,11]^. Besides, genes mediating cell density^[12]^, biofilm formation^[13]^, macromolecular repair^[14]^, and cell division^[15]^ have also been reported as critical for acid tolerance in *S. mutans*. To comprehensively elucidate its acid resistance mechanisms, studies at the system level have employed proteomic^[16]^ and transcriptomic^[17]^ profiling under varying pH and nutrient conditions, providing evidence and clues for the overall acid tolerance mechanism.

Although several key factors of acid tolerance have been well studied, our understanding of the roles that other genes play in this process still has notable gaps. Thus, genetic screening methods are required to elucidate the genetic basis for acid-tolerant phenotypes at the genome scale. The CRISPR-Cas system, a prokaryotic innate immune mechanism, confers acquired resistance against invasive genetic elements by cleaving targets (DNA or RNA) in a sequence-specific manner^[18]^. Recent advances in CRISPR-Cas programming have enabled genome-scale interrogation of gene function through transcriptional interference (CRISPRi)^[19]^. By synthesizing pooled single-guide RNAs (sgRNAs) via array-based oligonucleotide libraries, the Cas9 protein is directed to recognize target genes and suppress their expression. CRISPRi screens have been applied to study the impact of individual genes on bacterial fitness by quantifying sgRNA enrichment or depletion during bacterial growth using next-generation sequencing (NGS)^[20]^. Inspired by a previous study, which developed an arrayed CRISPRi library targeting 250 genes in *S. mutans*^[21]^, we developed a genome-wide pooled CRISPRi library targeting 95% predicted genes of *S. mutans*. This enabled genome-scale exploration of acid tolerance mechanisms in *S. mutans*, alongside functional and interaction analyses of the involved genes.

## Methods and Materials

### 2.1 Bacterial Strains, plasmid and growth condition

*S. mutans* UA159 was obtained from the American Type Culture Collection (ATCC, USA). The mutants and plasmid used in this study were listed in Table S1. The strains maintained at 20% glycerol stocks (- 80 °C) were plated on brain heart infusion (BHI) agar plates and cultured at 37 °C for 24 - 48 h (5% CO_2_). Single colony was selected to incubate in CDY (chemical defined medium (CDM) + 0.3% yeast extract (Oxoid, UK)) liquid medium for further study. CDYS (CDY with 1% (*w/v*) sucrose added) was used for subsequent biofilm-related assays.

### 2.2 Construction of *S. mutans* CRISPRi library

Based on the genomic information of *S. mutans* (NCBI RefSeq assembly GCF_000007465.2), sgRNAs were designed via the scripts developed by Bakker *et al.* (https://github.com/veeninglab/CRISPRi-seq). The design parameters were as follows: For each gene, one sgRNA was designed incorporating BbsI restriction enzyme sites, with the forward primer appended with the “ATGT” adhesive end sequence and the reverse primer appended with the “AAAC” adhesive end sequence. The protospacer adjacent motif (PAM) sequence was specified as “NGG”. A total of 1919 pairs of sgRNAs were synthesized (Beijing Ruibo Xingke Biotechnology Co., Ltd.).

The procedure for constructing a pooled sgRNA library was derived from a previous study^[22]^. Briefly, the synthetic single-stranded DNA oligonucleotides were annealed to form double-stranded DNA fragments. All annealed sgRNAs were mixed in an equimolar ratio to form an sgRNA library. The sgRNA library was then phosphorylated using TPNK (NEB M0525S). To express the sgRNA in *S. mutans*, we constructed a suicide plasmid pYL02 (Supplementary materials for sequence), which carried an ampicillin resistance gene and an origin of replication (ori) enabling its replication and antibiotic selection in *Escherichia coli* (*E. coli*). It additionally contained an erythromycin resistance and sgRNA expression cassette. Within this cassette, sgRNA expression was driven by the P*_3_*promoter, while erythromycin resistance was expressed under the control of the P*_23_* promoter. Two homology arms flanked this cassette, enabling its targeted integration into specific genomic loci via homologous recombination. The pYL02 was digested with the restriction enzyme BbsI. The linearized vector was then ligated to the phosphorylated sgRNA pool using T4 ligase.

The ligation products were collected and then electro-transformed into *E. coli* stbl3 competent cells (Transgene). The cells were plated onto LB agar plates containing ampicillin (100 μg/mL) using sterile glass beads. After an overnight incubation, 2.1 × 10^6^ colonies from the plate were collected into LB media and mixed thoroughly. Subsequently, 5 mL of the bacterial suspension was mixed with an equal volume of 50% glycerol (*v/v*) and stored at -80 °C. The remaining bacterial suspension was used for plasmid extraction with a Plasmid Maxi Kit (Qiagen). Next-generation sequencing (NGS) was used to verify the presence and distribution of sgRNA within the pool. Briefly, the sgRNA region was amplified using the high-fidelity DNA polymerase Phanta (Vazyme, Beijing). The PCR products were purified and then quantified using the Qubit dsDNA HS Assay Kit. High-throughput sequencing was performed using the Illumina MiniSeq. DESeq2 was used to analyze the processed data to calculate the abundance of each sgRNA.

To achieve the *S. mutans* with xylose-inducible dspCas9, we deleted the endogenous *cas9* via in-frame deletion. We integrated an exogenous copy of dspCas9 controlled by the TC-Xyl inducible cassette into the *phnA-mtlA* locus through overlapping PCR and homologous recombination as previously described^[23,24]^.

The constructed sgRNA plasmid pool was transformed into the above strain via CSP-mediated transformation^[23]^, and the constitutively expressed sgRNA was integrated into the *phnA-mtlA* gene region. Transformants were selected by BHI plates containing 12 μg/mL erythromycin (Figure 1A).

**Figure 1.**
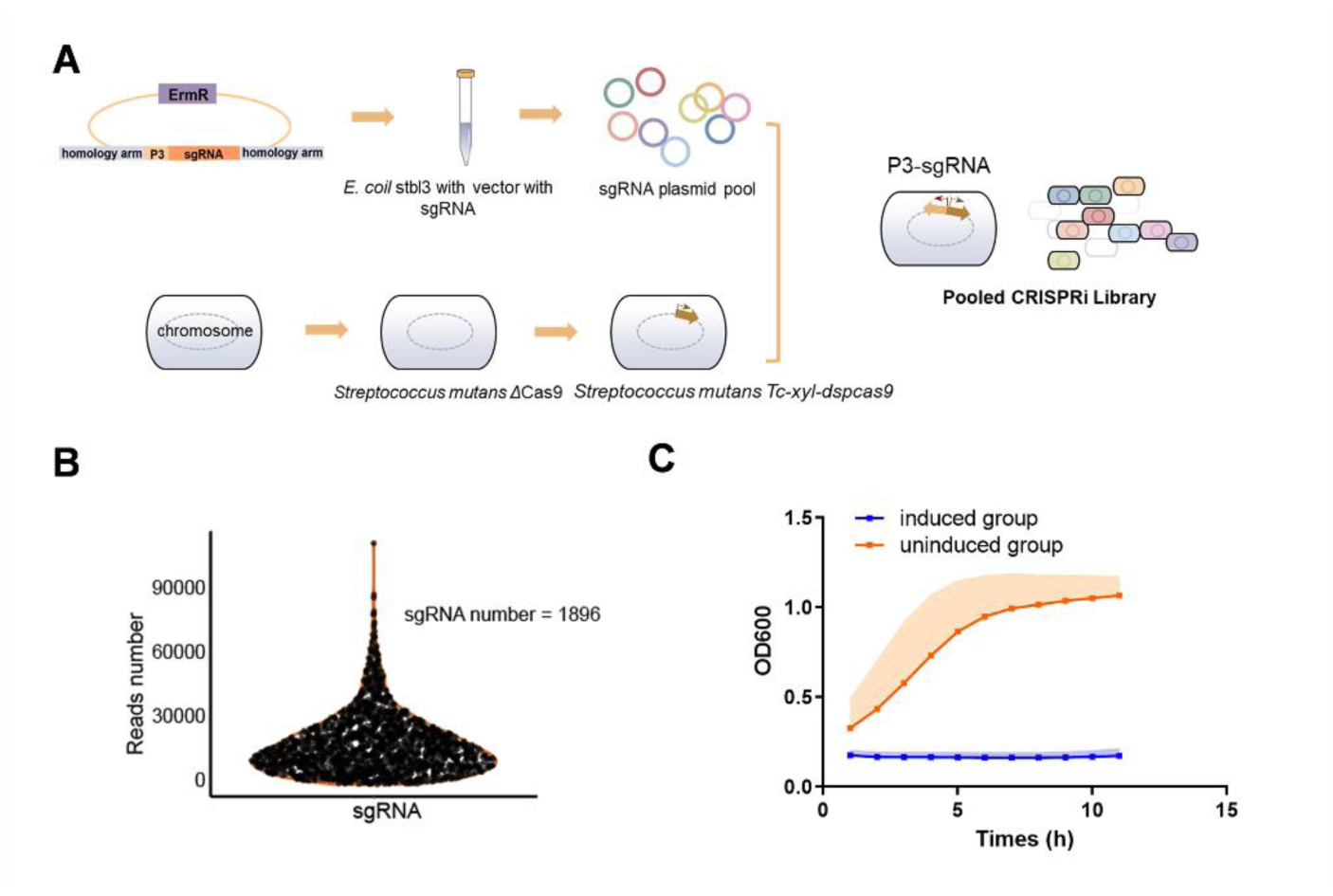
Construction and validation of CRISPRi library. (A) Schematic overview for CRISPRi library construction of *S. mutans*. We designed 1919 sgRNAs for the genome of *S. mutans*. The plasmid vector pYL02 was digested and then ligated to the phosphorylated sgRNA pool. The ligation products were collected and then electro-transformed into *E. coli* stbl3 competent cell to enrich the plasmid. The sgRNA plasmid pool was harvested from the successful transformants. We deleted the endogenous *cas9* via in-frame deletion and the TC-Xyl inducible dspCas9 were integrated into the chromosome. The constructed sgRNA plasmid pool was transformed into the above strain via CSP-mediated transformation. (B) NGS of the pool revealed that the sgRNA counts were approximately normally distributed. (C) Growth curve of FtsZ CRISPRi strain under 1% xylose induced or uninduced conditions.

### 2.3 CRISPRi-seq

#### 2.3.1 Sample preparation of CRISPRi-seq

We used the genome-wide sgRNA library to perform screening experiments. Briefly, the library was resuscitated in CDY medium at a 1:20 dilution ratio and cultured until reaching the logarithmic growth phase. Subsequently, the library was inoculated into five different media: (1) THY (Todd-Hewitt Broth+0.3% yeast extract) media (pH 7.4) (2) THY media supplemented with 1% xylose (*w/v*) (pH 7.4) (3) CDY media (pH 7.4) (4) CDY media supplemented with 1% xylose (*w/v*) (pH 7.4) (5) CDY media supplemented with 1% (*w/v*) xylose (pH 5.0) and cultured until reaching the logarithmic growth phase. This procedure was repeated three times, and the samples were collected. By comparing groups (1) and (2), (3) and (4), we identified essential genes of *S. mutans* growth in different media, while the acid-tolerance related genes were assessed through the gene abundance comparison of groups (4) and (5) (Figure 2A).

**Figure 2.**
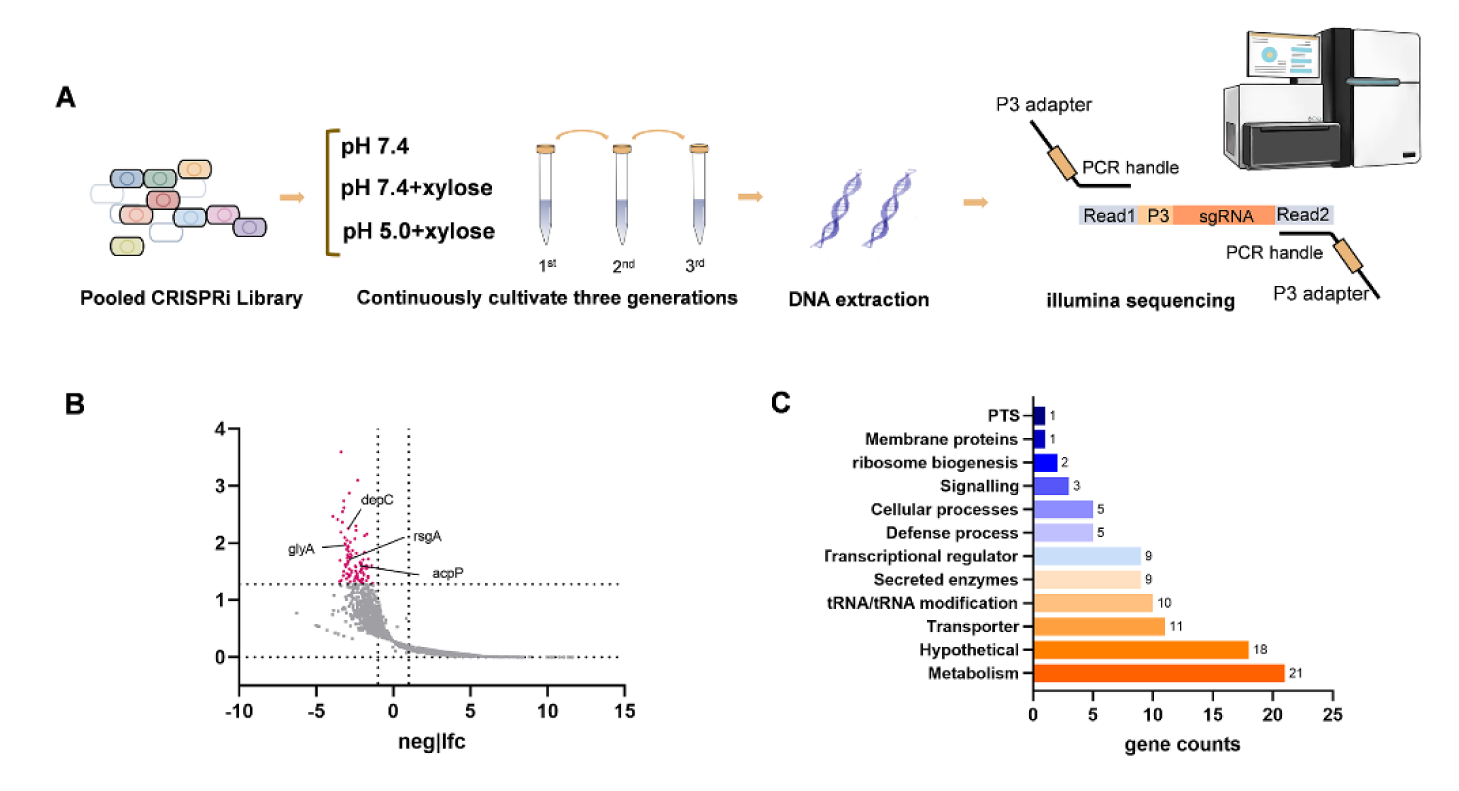
Genome-scale CRISPRi screening revealing essential genes and acid-tolerant genes. (A) Schematic overview for CRISPRi screening in *S. mutans*. NGS was used to determine the changes in sgRNA abundance. (B) Volcano plot of acid-tolerant genes. NGS sgRNA abundance comparison of CRISPRi library exposed to acid condition (pH=5.0, 1% xylose) or to the control condition (pH=7.4, 1% xylose) (*n* = 3 for each condition). Genes with absolute |fold change| > 2.0 and an adjusted *P*-value < 0.05 were considered to be acid-tolerant genes. Genes were functionally categorized using annotations from the Kyoto Encyclopedia of Genes and Genomes (KEGG) and Gene Ontology (GO) databases. (C) Quantification of the number of acid-tolerant genes for different functional categories.

The total DNA was extracted using the TIANamp Bacteria DNA kit (Tiangen, China). Concentration and quality measurements of the extracted DNA were performed with a NanoDrop ND-1000 spectrophotometer (Thermo Fisher Scientific, Waltham, MA, USA).

#### 2.3.2 sgRNA sequencing and data analysis

DNA samples collected from the CRISPRi screen were used to construct a sgRNA library through a two-step PCR procedure^[25]^. In the initial PCR step, the cassette containing the sgRNA sequence was amplified. The amplified products were then processed in a secondary PCR using a primer pair that encoded a unique 8-bp barcode, required for multiplexing, along with a stagger sequence to increase library complexity, as described^[25]^. The primers and PCR procedure were listed in the Supplementary materials. Products of the second PCR were purified, pooled, diluted, and mixed with 10% PhiX, and then sequenced using the Novaseq 6000 (Illumina). The low-quality sequencing reads were first removed using the fastp program 0.20.0, as described in a previous study^[26]^. The trimmed reads were then used to map sgRNA sequences to the S. mutans library using MaGeCK (Version 0.5.9.5). Read counts of sgRNA for each sample were quantified by MAGeCK^[27]^. Count data were filtered, normalized, and ranked by MAGeCK^[27]^. The reads observed from the induced group were divided by the uninduced reads to obtain a ratio. For any given gene, a ratio ranging from 0.01 to 0.1 suggested compromised fitness, while a ratio below 0.01 signified essentiality of the gene^[28]^. Acid-tolerance genes were identified using a threshold of |fold change| > 2.0 and an adjusted *P*-value < 0.05. Functional categorization^[29]^ of candidate genes was performed based on the enrichment analysis and annotations from the Kyoto Encyclopedia of Genes and Genomes (KEGG) and Gene Ontology (GO) databases. We analyzed protein-protein interactions (PPIs) using the STRING database to elucidate the functional associations between candidate genes and analyzed them using Cytoscape (version 3.10.3).

### 2.4 Construction of *S. mutans* mutants

This study constructed the *glyA*, *deoC*, *acpP*, and *rsgA* null-mutant strains of *S. mutans* UA159 using the IFDC2 cassette through overlapping PCR and homologous recombination, as described in previous studies^[23, 24]^. The primers used in this study were listed in Table S2.

### 2.5 Growth curve assay

Bacterial growth of wild-type strain (WT, UA159), *ΔglyA*, *ΔdeoC*, *ΔacpP*, and *ΔrsgA* was quantified using a microplate reader (SpectraMax 190, Molecular Devices, USA). Overnight cultures were diluted 1:100 in CDY medium (pH 5.0), with uninoculated medium serving as the control. The results of optical density (OD600) were recorded every 30 min for 24 h using Molecular Devices software, with 5 s shaking before each measurement. Growth curves were derived from time-dependent OD600 values.

### 2.6 Acid tolerance assays

Acid tolerance assays were conducted as previous study described with slight modifications^[30]^. To assess the acid tolerance of *S. mutans* and its four mutants, we collected the bacterial suspension at the logarithmic growth phase (OD600 = 0.5) by centrifugation at 4,000 g for 10 min at 4 °C and washed it once with 0.1 M glycine buffer (pH 7.0). A portion was directly diluted by PBS, plated on BHI agar, and incubated at 37°C under 5% CO_2_ for 48 h. The Colony-forming units (CFU) before the treatment served as the control.

1. A portion of the culture was directly exposed to 0.1 M glycine-HCl (pH 2.8). The bacterial suspensions were mixed thoroughly by vortex and incubated at pH 2.8 for 10 min.
2. To induce the acid tolerance response (ATR), another portion was resuspended in CDY at pH 5.0 and incubated for 2 h before being transferred to 0.1 M glycine buffer (pH 2.8). The bacterial suspensions were mixed thoroughly by vortexing and incubated at pH 2.8 for 10 min.

Surviving cells were then diluted with PBS, plated on BHI agar, and incubated at 37°C under 5% CO_2_ for 48 h. The CFU was calculated from the plate. The viability rate was calculated according to the formula:

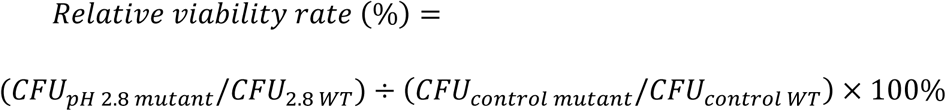

### 2.7 Biofilm formation and confocal laser scanning microscopy (CLSM) observation

The bacterial suspension of WT and four mutants in the logarithmic growth phase (OD=0.5) was diluted 100 times by CDYS medium and incubated anaerobically at 37 °C for 48 h. Biofilm images were captured using CLSM with a 60×oil immersion objective lens. (TCS-SP8 and MICA, Leica). Imaging gates were set to 488 nm for SYTO 9 stain and 561 nm for propidium iodide. Each biofilm was scanned at 3 random positions.

### 2.8 Statistical analysis

Statistical analysis was conducted using GraphPad Prism 7 software (version 7.00 for Windows; GraphPad Prism, Inc., La Jolla, USA). After conducting a homogeneity of variance test (Levene’s test) on the data, one-way ANOVA and the SNK method were used for multiple comparisons between the groups. Both groups were subjected to a double-sided test, with a significance level of α= 0.05. All figures were generated with GraphPad Prism 7 software.

### 2.9 Data availability

The sgRNA sequencing data have been submitted to the public database Sequence Read Archive with accession number PRJNA1285880 https://dataview.ncbi.nlm.nih.gov/object/PRJNA1285880?reviewer=9fluac1b7llnn52khcjbsslt9f. All datasets produced and/or examined in this study can be obtained from the corresponding author upon reasonable request.

## Results

### 3.1 Construction of a genome-wide sgRNA library for *S. mutans*

First, we created an sgRNA library to repress each annotated gene in *S. mutans* (Figure 1A). Oligonucleotides for a total of 1919 sgRNA sequences were synthesized, ligated to the plasmid vector pYL02, and cloned into *E. coli* Stbl3 competent cells. 2.1 × 10^6^ colonies were collected to ensure at least 1000-fold coverage. Next-generation sequencing (NGS) was used to verify the presence and distribution of sgRNA within the pool. The results showed that 98.8% of sgRNAs (1896 sgRNAs) were represented with at least one sequencing read in NGS analysis, and the NGS accuracy was 94.63%. DESeq2 was used to analyze the processed data to calculate the abundance of each sgRNA. The DESeq function was utilized for differential analysis to quantify the abundance changes of each sgRNA. The distribution of sgRNAs was shown in Figure 1B, which showed approximately normal distribution. The plasmid pool was transformed into *S. mutans*, and 2 × 10^4^ colonies were obtained, 100-fold coverage. To evaluate the efficiency of CRISPRi in reducing gene expression, we focused on *ftsZ*, which encodes the conserved tubulin-like cell division protein FtsZ, playing a crucial role in cell division. As illustrated in Figure 1C, xylose-induced (1%) activation of dead SpCas9 (dCas9) significantly impaired cellular growth in strains containing *ftsZ*-targeting sgRNAs, whereas the non-induced counterparts maintained normal proliferation rates.

### 3.2 Determination of essential genes of *S. mutans*

CRISPRi screen with xylose-inducible library was performed in CDY medium and THY medium, and the procedure was shown in Figure 2A. The sgRNAs were amplified by two-step PCR and subsequently quantified by Illumina sequencing. Sequencing verified the presence of all 1896 sgRNAs in the uninduced sample of the xylose-inducible library. The determination criteria of essential genes were consistent with the previous report by Shields RC et al.^[28]^. For a specific gene, a ratio ranging from 0.01 to 0.1 suggested compromised fitness, while a ratio below 0.01 indicated the gene’s essentiality. In this study, we found 152 essential genes. 72.4% of the essential genes identified through our CRISPRi-seq method have also been recognized as essential (57.9%) or compromised (14.5%) in previous Tn-seq studies, demonstrating a high level of consistency between these methodologies^[21, 28]^ (Table S3).

### 3.3 Identification and functional analysis of acid tolerance-related genes

To identify acid resistance-related genes, we performed CRISPRi sequencing to compare sgRNA abundance under normal (pH 7.4, CDY medium +1% xylose) and acidic conditions (pH 5.0, CDY medium +1% xylose) (Figure 2A), through which we identified 95 depleted candidate genes (Figure 2B). To better understand these genes, we categorized them based on their functional annotations (Figure 2C and Table S4). The most abundant category within the depleted genes was metabolic processes (21 genes), including the acid tolerance-confirmed gene *plsX*. Major functional groups consisted of cofactor biosynthesis (6 genes) and amino/nucleotide sugar metabolism (4 genes). It is noteworthy that our CRISPRi screen revealed significant depletion of tRNA/tRNA modification-related sgRNAs under acidic conditions, including six tRNA genes and four tRNA modification genes, indicating the essential role of tRNA metabolism in cellular survival under acidic stress (Table S4). The transcriptional regulators also demonstrated critical roles in acid tolerance, with 4 out of 9 identified genes (*brpA*, *ccpA*, *clpP*, and *sloR*) having been experimentally validated. Furthermore, key signaling components (e.g., *vicK* and *gcrR*), defense systems (including *cas2* and *cas5* of the CRISPR-Cas apparatus), and multiple transporters were also involved in acid adaptation (Table S4).

### 3.4 Interactions between the acid-tolerance related genes

To explore the connections between 95 depleted candidate genes, we constructed a PPI network using the STRING database. We found that out of the 95 depleted candidate genes, 57 genes exhibited interaction relationships (Figure 3). The constructed PPI network exhibited scale-free properties with a clustering coefficient of 0.372 and the enrichment *P*-value of 0.0385, suggesting biological relevance beyond random interactions. We further analyzed the hub proteins based on degree centrality analysis (Table S5) and MCC algorithm (Table S6) using cytoscape, which revealed *comYC*, *SMU_1979c*, *deoC*, *acpP*, *nadD,* and *SMU_1988c* encode the hub proteins. DeoC and AcpP have been demonstrated to be associated with acid tolerance in the aforementioned experiments. Notably, the acyl carrier protein AcpP (degree = 5) and nicotinate-nucleotide adenylyltransferase NadD (degree = 5) showed extensive cross-modular connections, positioning them as a potential regulatory nexus in the acid stress response. Network modules were identified using MCODE with parameters set as: degree cutoff = 2, node score cutoff = 0.2, k-core = 2, max depth = 100. We identified two functional modules through MCODE analysis: the first module (M1), containing ComYA, SMU_1988c, SMU_1979c, and ComYC, was identified as a type II protein secretion system complex, while the second module (M2), comprising DeoC, SMU_1118c, SMU_1121c, and Cdd, showed significant association with the pyrimidine nucleoside metabolic process, ABC transporters, and permease. The two modules may play a significant role in acid tolerance, and further study is needed to confirm.

**Figure 3.**
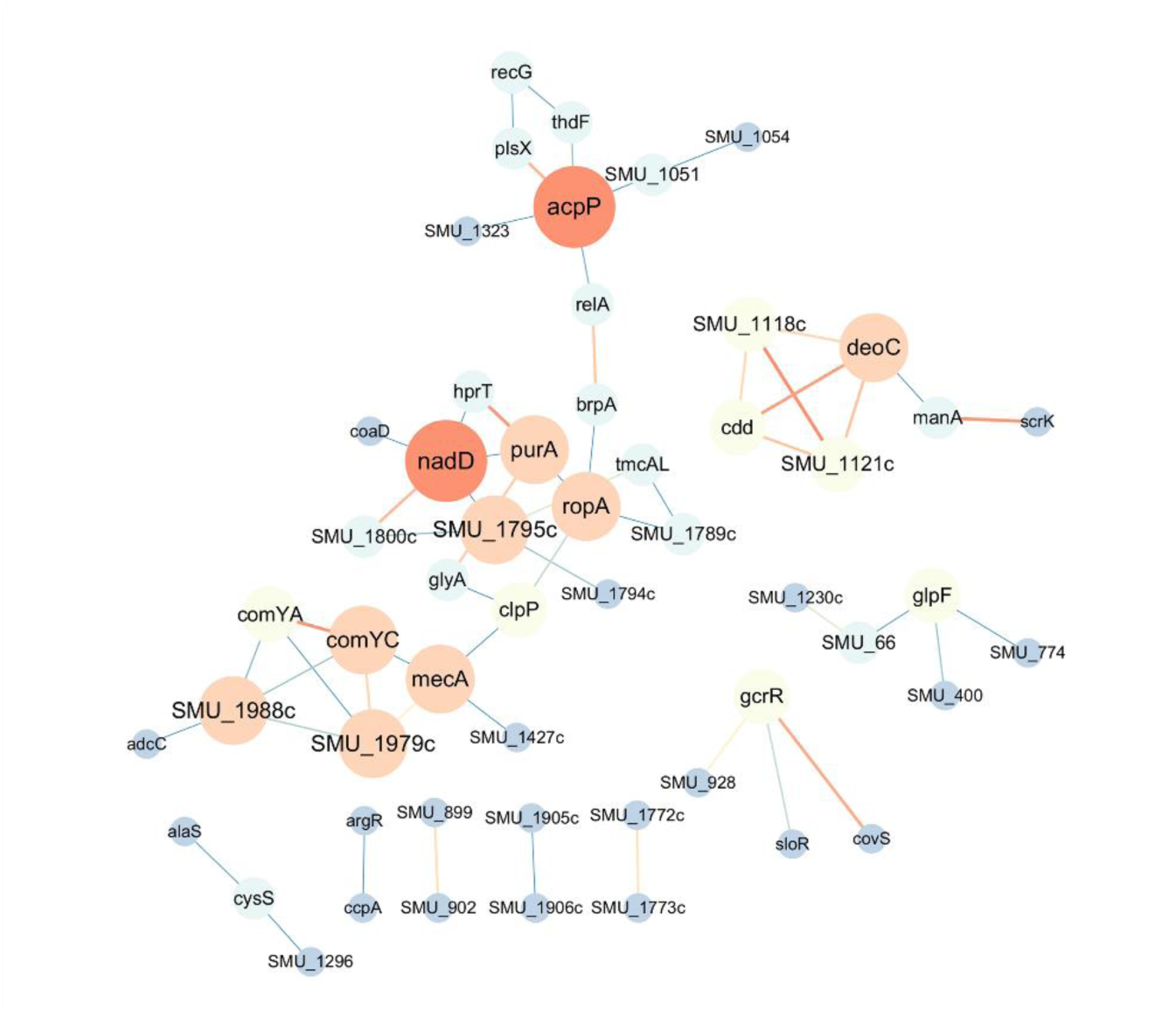
PPI network of 95 acid-tolerant genes generated by the STRING database and analyzed using Cytoscape. Node size was proportional to the number of interaction edges (degree), with larger, more orange nodes indicating higher connectivity and smaller, more blue nodes indicating lower connectivity. Similarly, edge thickness corresponded to the interaction strength between nodes, where thicker, darker orange edges represented stronger interactions, while thinner, paler blue edges represented weaker interactions.

### 3.5 Validation of the acid-tolerance related genes

To further validate the sequencing results, we constructed null mutants for several genes identified in the results. Subsequently, a series of phenotype experiments were carried out with both WT and the generated null mutants to investigate the impact of gene deletion on their acid tolerance. Compared with WT, the growth of mutants was suppressed in the acidic condition (pH 5.0), mainly manifested as the delayed entry into the logarithmic growth phase and reduced total bacterial counts during the stationary phase (Figure 4A). Two acid tolerance assays were used to evaluate the constitutive tolerance (Figure 4B) and acid-induced tolerance (Figure 4C) of *S. mutans*. *ΔglyA*, *ΔdeoC*, *ΔacpP*, and *ΔrsgA* exhibited a significantly lower survival rate than WT after 10 min of lethal acid shock in two assays (*P* < 0.05, Figure 4B and 4C), which indicated the mutants were more sensitive to acid conditions. Interestingly, *ΔglyA* and *ΔacpP* showed lower viability after the pH 5.0 adaptation, while *ΔdeoC* and *ΔrsgA* showed higher viability, which may indicate that these genes play different roles in acid tolerance. We also measured the 48 h mature biofilms using CLSM and found that the biofilms of the 4 mutants contained more dead cells compared to the WT (Figure 4D and 4E), suggesting an impaired acid tolerance ability in the mutants.

**Figure 4.**
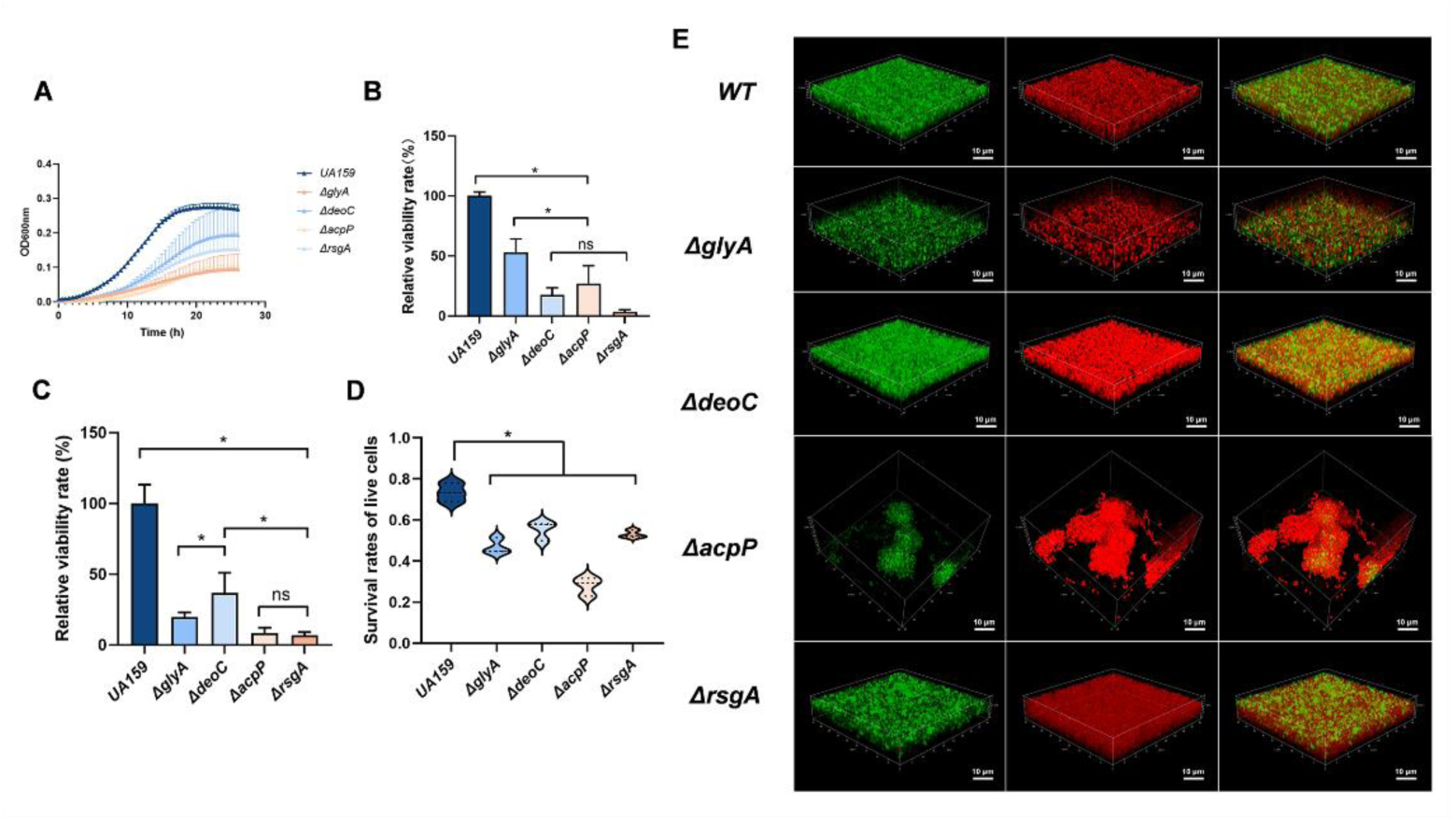
Validation of the acid-tolerance related genes. (A) Growth curves of UA159, *ΔglyA*, *ΔdeoC*, *ΔacpP*, and *ΔrsgA* strains at the acid condition (pH=5.0). (B) Relative viability rate of UA159, *ΔglyA*, *ΔdeoC*, *ΔacpP*, and *ΔrsgA* strains in direct acid kill assay. (C) Relative viability rate of UA159, *ΔglyA*, *ΔdeoC*, *ΔacpP*, and *ΔrsgA* strains in acid-tolerant response assay. (D)Survival rates of live cells of WT and mutants. (E) CLSM images of UA159, *ΔglyA*, *ΔdeoC*, *ΔacpP*, and *ΔrsgA* strains 48 h biofilm. The green fluorescence indicates live cells, while the red fluorescence indicates dead cells. Images were examined at 60× objective magnification. Scale bar: 10 µm (*, *P* < 0.05; ns, not significant).

## Discussion

*S. mutans* is recognized as the primary pathogen in cariogenesis, with acid tolerance constituting a crucial virulence factor. Although previous investigations have employed sequencing approaches to explore its acid-tolerant mechanisms and characterized several classical pathways, there remains a paucity of genome-wide comprehensive studies. In our study, we reported the first construction of a genome-wide CRISPRi library for *S. mutans*. Through next-generation sequencing analysis, we systematically profiled essential genes and acid tolerance-associated genes. Our findings also identified the hub acid tolerance-related genes and functional modules, providing novel insights into the genomic basis of acid adaptation mechanisms in *S. mutans*.

The identification of essential genes in *S. mutans* is of significant importance for understanding its fundamental biological processes and discovering novel antimicrobial targets. Previous studies employing Tn-seq analysis identified 203 essential genes in blood agar and 319 in FMC medium. Subsequently, the team constructed an arrayed CRISPRi library comprising 250 genes targeting the essential genes identified above. Validation demonstrated that 162 of these genes were essential. Similarly, in this study, we identified 152 essential genes in THY media and 255 essential genes in CDY media, demonstrating condition-dependent variations in essential gene profiles. Comparison of THY and Tn-seq data revealed that among the 152 essential genes identified, 136 were covered by Tn-seq. Of these, 88 overlapped with previously reported essential genes, while 22 matched compromised genes and 25 corresponded to non-essential genes, demonstrating strong concordance between the methods. The remaining 16 essential genes were not detected by Tn-seq^[28]^. We propose two possibilities underlying these differences. Firstly, the use of different culture media between studies may influence essential gene selection. Secondly, the inherent limitations of the Tn-seq methodology contribute to potential misclassification, including insertion biases of transposon systems due to sequence-specific insertion preferences, the lack of transposon insertion sites, and the lack of transposon insertion density^[28]^. Similarly, CRISPRi also had limitations. CRISPRi may fail to detect essentiality due to suboptimal sgRNA design or suppressor mutations that bypass essentiality^[21]^. The gene silencing effects of CRISPRi are lower than those of Tn-seq, and inducer concentration may influence the efficacy of gene silencing^[31]^.

We employed the CRISPRi-based library to investigate acid-tolerant associated genes in *S. mutans*. A key advantage of this approach lay in its capacity to systematically interrogate all genes, including those classified as essential under conventional transposon mutagenesis^[32]^. However, our analysis revealed unexpected results: classical acid tolerance pathways, such as F-ATPase, fatty acid biosynthesis networks, and agmatine deiminase system, were conspicuously absent from the enriched genes. We analyzed the results for these key genes and observed significant abundance depletion under acid stress conditions for the majority (Table S7). Subsequently, we also identified genes that have been confirmed to be acid-tolerance-related in these systems: *fabM*^[17]^*, fabD*^[33]^*, and aguR*^[34]^. Consistent with these reports, these genes were also significantly depleted in our study. Furthermore, it has been reported that deletion of the *atpF* gene did not significantly impair intracellular pH maintenance capability. This indicates that not all genes within the systems mentioned above are essential for *S. mutans* acid tolerance, suggesting that potential compensatory mechanisms may exist. Additionally, within these classical pathways, a substantial proportion of genes (such as *fabG* and *atpD*) exhibited growth restriction under induction conditions (Table S7). The impact of growth restriction manifested as an excessively low abundance of this gene under induction conditions. Despite a further reduction in gene abundance under acidic conditions, these changes were overlooked in statistical analyses due to the presence of low-count bias. Paradoxically, this limitation conferred a unique advantage. The screened genes demonstrated a direct link to acid tolerance, as their silencing specifically compromised acid survival without affecting baseline cellular proliferation. In this study, we observed significant depletion of tRNAs and tRNA modification-associated genes under acidic conditions. As essential intermediaries in protein synthesis, tRNAs play a pivotal role in translation by mediating interactions between mRNA codons and amino acids. This fundamental role requires their specific engagement with multiple molecular partners, such as ribosomes, elongation factors, mRNAs, and aminoacyl-tRNA synthetases^[35]^. Experimental studies have demonstrated that defects in tRNA modifications impair bacterial resistance to environmental stresses, which have been validated in *Deinococcus radiodurans*^[36]^ and *Streptococcus thermophilus*^[37]^. In *S. mutans*, the t^6^A-deficient mutants are compromised in biofilm formation and become more sensitive to low pH and oxidative stress than the wild-type^[38, 39]^. Moreover, aminoacyl-tRNA synthetases (aaRSs) ensure translational fidelity through their proofreading activity. Loss of these enzymes may compromise the accuracy of bacterial protein synthesis. However, emerging evidence suggests that mistranslation may paradoxically enhance environmental adaptability by generating proteomic diversity under stress conditions^[40]^. Hence, further research is still needed to explore the role of tRNA and its modifications in acid tolerance.

Our subsequent PPI network analysis revealed AcpP (encoding acyl carrier protein) and NadD (nicotinamide-nucleotide adenylyltransferase) as hub proteins with the highest connectivity. Notably, acyl carrier protein—a key protein in fatty acid metabolism—has been previously demonstrated to play a critical role in acid tolerance in *Pediococcus acidilactici*^[41]^. In this study, we experimentally validated the role of AcpP in acid tolerance in *S. mutans* by constructing an *acpP* null mutant. Our findings are consistent with previous studies, thereby reinforcing the general importance of AcpP in acid tolerance mechanisms. Nicotinamide adenine dinucleotide (NAD) and its reduced form NADH serve as common cofactors for many oxidoreductases that participate in diverse metabolic pathways^[42]^. Within the metabolic framework, NadD-family NaMN adenylyltransferases (NaMNATs) catalyze the conversion of NaMN precursors to NAD, which has been extensively investigated as an antimicrobial target ^[43]^. Notably, no previous studies have systematically evaluated the role of NadD in bacterial acid tolerance. We postulated that NadD modulated acid tolerance primarily through its regulatory influences on metabolism, particularly in pathways governing redox homeostasis and energy production^[44]^. Furthermore, we identified two functionally distinct gene modules. Module M1 comprised *comYA*, *SMU_1988c*, *SMU_1979c*, and *comYC*, where *comYA* and *comYC* have been experimentally validated to mediate DNA uptake during bacterial competence^[45, 46]^. Module M2 included *deoC*, *SMU_1118c*, *SMU_1121c*, and *cdd*. Notably, *deoC* has been implicated in biofilm dispersal-mediated infection dissemination and neutrophil evasion through NET degradation^[47]^. However, no studies on acid tolerance-related aspects of these modules have been conducted to date. Our data preliminary suggested its potential involvement in acid tolerance; however, both modules necessitate systematic investigations to conclusively understand their roles in acid tolerance mechanisms.

In summary, this study successfully constructed the first genome-wide CRISPRi library for *S. mutans* and applied it to screen essential genes and acid tolerance-related genes. Through functional analysis and PPI network analysis, we identified pathways, genes, and functional modules linked to acid tolerance. These findings provide a comprehensive understanding of the acid tolerance mechanisms of *S. mutans*.

## CRediT authorship contribution statement

Yaqi Chi: Methodology, Investigation, Formal analysis, Writing the original draft. Yuxing Chen: Methodology, Investigation, Formal analysis, Writing the original draft. Chongyang Yuan: Methodology, Investigation, Formal analysis. Liuchang Yang: Investigation. Mingrui Zhang: Investigation. Xiaolin Chen: Investigation. Yiran Zhao: Investigation. Ming Li: Writing-review and editing, Supervision. Xiaoyan Wang: Resources, Funding acquisition, Project administration. Yongliang Li: Methodology, Investigation, Validation, Formal analysis, Conceptualization, Writing-review and editing, Supervision.

## Acknowledgements

This work was supported by the Beijing Natural Science Foundation: 7222220, Research Foundation of Peking University School and Hospital of Stomatology: PKUSS20230117, National Natural Science Foundation of China (82001039), the Fundamental Research Funds for the Central Universities and Young Elite Scientist Sponsorship Program by CAST (No. 2019QNRC001 to YL.L). We appreciate the help of Dr. Jiao Liu from the Center of Medical and Health Analysis, Peking University Health Science Center, in confocal microscopic imaging.

